# Identification of Discriminative Gene-level and Protein-level Features Associated with Gain-of-Function and Loss-of-Function Mutations

**DOI:** 10.1101/2021.01.01.424981

**Authors:** Cigdem S. Bayrak, Aayushee Jain, David Stein, Kumardeep Chaudhary, Girish N. Nadkarni, Tielman Van Vleck, Anne Puel, Stephanie Boisson-Dupuis, Satoshi Okada, Peter D. Stenson, David N. Cooper, Avner Schlessinger, Yuval Itan

## Abstract

Identifying whether a given genetic mutation results in a gene product with increased (gain-of-function; GOF) or diminished (loss-of-function; LOF) activity is an important step toward understanding disease mechanisms as they may result in markedly different clinical phenotypes. Here, we generated the first extensive database of all currently known germline GOF and LOF pathogenic mutations by employing natural language processing (NLP) on the available abstracts in the *Human Gene Mutation Database*. We then investigated various gene- and protein-level features of GOF and LOF mutations by applying machine learning and statistical analyses to identify discriminative features. We found that GOF mutations were enriched in essential genes, autosomal dominant inheritance, protein binding and interaction domains, whereas LOF mutations were enriched in singleton genes, protein-truncating variants, and protein core regions. We developed a user-friendly web-based interface that enables the extraction of selected subsets from the GOF/LOF database by a comprehensive set of annotated features, and downloading up-to-date versions (https://itanlab.shinyapps.io/goflof/). These results could ultimately improve our understanding of how mutations affect gene/protein function thereby guiding future treatment options.

## Introduction

To understand the mechanisms underlying human genetic diseases, it is crucial to study the effects of genetic mutations on their protein products. Based on their effect on protein function, mutations can be categorized into two main types: gain-of-function (GOF) mutations, which enhance the activity of the mutated protein, and loss-of-function (LOF) mutations which cause partial or complete protein inactivation ^1^. Over the past decades, it has been shown that GOF and LOF mutations in the same gene may result in completely different disease phenotypes ^1–6^. For example, GOF mutations in the *STAT1* gene cause chronic mucocutaneous candidiasis (CMC), which is characterized by recurrent disease of the nails, skin, and oral and genital mucosae, whereas dominant LOF mutations in *STAT1* cause Mendelian susceptibility to mycobacterial disease (MSMD), which is characterized by predisposition to clinical disease caused by weakly virulent mycobacteria ^7, 8^. *WAS*, *MDA5*, *PCSK9*, and *CACNA1C* are additional known examples of genes in which GOF and LOF mutations impact different genetic pathways, leading to distinct clinical phenotypes that require different drug treatment regimens ^2–6^. To determine the disease phenotype that can in turn guide the identification of drug targets, it is essential to understand the different functional effects of mutations, i.e., whether the protein enhances or diminishes activity.

Whether a specific mutation causes a loss- or gain-of-function is ultimately determined by its impact on the biological pathway(s) associated with the protein(s) encoded by the mutated gene^9^. Mutations can have a significant effect on protein stability, hydrogen bond network, conformational dynamics, as well as on binding to other molecules, and many other important biochemical and physiological properties of proteins ^9–11^. Thus, to understand the genetic mutation’s effect on protein structure and function, it is necessary to accurately and reliably distinguish between GOF and LOF mutations.

Understanding the properties of genetic variants identified by of next-generation sequencing (NGS) is of great interest for human health and genetic research ^12–14^. So far, various computational deleteriousness and pathogenicity prediction tools based on machine learning algorithms and DNA sequence information, such as CADD, PolyPhen-2, SIFT and REVEL, have been developed to facilitate the identification of disease-causing mutations ^15–18^. For example, CADD integrates multiple annotations including functional information and conservation, whereas SIFT and PolyPhen-2 mainly consider sequence conservation and biochemical features, whilst REVEL integrates scores from multiple prediction tools. However, none of these tools have the capability to distinguish GOF from LOF mutations.

To date, there has only been a limited effort to identify and analyze the discriminative features of GOF and LOF mutations, and these studies have numerous limitations. Thus, no study to date has attempted to generate a comprehensive public germline GOF/LOF database that could prove invaluable for developing an accurate and generalizable GOF/LOF classifier and for annotating NGS variants. For example, in a recent study, a computational method to predict missense GOF and LOF variants in voltage-gated sodium and calcium channels, *SCNxA* and *CACNA1x* family genes, has been presented ^19^. In a separate study, 129 GOF mutations from 59 genes and 258 LOF mutations from 109 genes were analyzed and six features were proposed to be discriminant^20^. Additionally, a computational method, HMMvar-func, that predicts functional outcome of a mutation has been reported ^21^. HMMvar-func was however trained only on two genes, *TSHR* and *TP53*, and it defines GOF mutations as having acquired a new function, without including GOF mutations with enhanced activation, which accounts for the majority of GOF mutations ^22–24^.

Other shortcomings of such existing studies include: (i) the GOF and LOF mutations were not shown to be disease-causing; (ii) the number of incorporated features was limited; and (iii) they lacked publicly accessible data. In particular, no study has generated a comprehensive database of GOF and LOF mutations.

In this study, we applied various machine learning and statistical approaches to construct a comprehensive database of currently known disease-causing GOF and LOF mutations from the Human Gene Mutation Database (HGMD) ^25^. We first applied text-mining and natural language processing (NLP) to abstracts of published pathogenic mutations to extract GOF and LOF mutations from HGMD (Professional version 2019.1), and then annotated 12 protein-level, 11 gene-level, and 10 variant-level features that were broken down into 1,725 variables. We demonstrated that there are variant-, gene-, and protein-level features that serve to distinguish GOF and LOF mutations, and discuss the biology of these discriminant features (Fig. 1). Finally, we generated and made publicly available the first extensive database of 9,541 pathogenic GOF and LOF mutations based on the HGMD December 2019 release (https://itanlab.shinyapps.io/goflof/). The purpose of the web-based interface is to provide a publicly available GOF/LOF database for the scientific community to analyze, extract, and download the available mutations as well as a comprehensive set of variant-, gene-, and protein-level features.

**Fig. 1.**
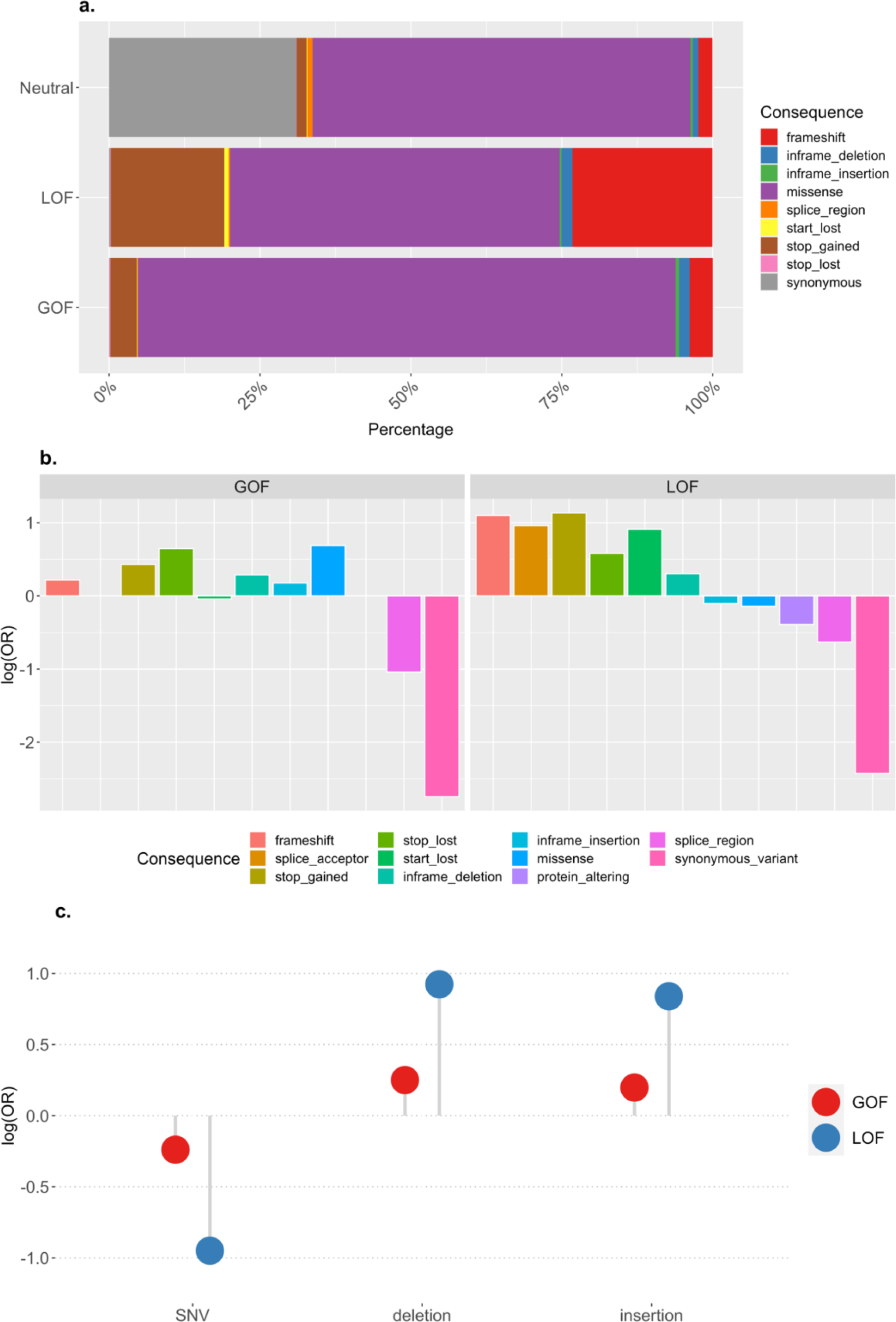
Variant consequence and class types. **a)** Predicted mutational consequence percentages for each mutation type, including GOF, LOF and likely neutral variants. **b)** Predicted mutational consequences of GOF/LOF mutations. The y-axis corresponds to the log10 of the odds ratio of the enrichment of predicted consequences in GOF and LOF mutations compared to likely neutral variants. **c)** Predicted variant classes. The y-axis corresponds to the log10 of the odds ratio of the enrichment of variant classes in GOF and LOF mutations compared to likely neutral variants.

Throughout this paper we use the term ‘mutation’ to refer to pathogenic variants.

## Methods

### Construction of GOF and LOF database

#### Selection of GOF and LOF mutations using NLP

We used the HGMD database (Professional version, 2019.1 release) that contains 256,070 mutations of which 202,639 were from the disease-causing class (DM, highest confidence for being pathogenic) ^25, 26^. For each disease-causing mutation, we searched all associated paper titles and abstracts for possible acronyms of GOF and LOF mutations including “gain of function(s)”, “gain-of-function(s)”, “GOF”, “loss of function(s)”, “loss-of-function(s)”, “LOF” by using phrase-based matching in spaCy v2.1.3 open-source natural language processing (NLP) library in Python 3.7.3 (https://spacy.io). The NLP text processing pipeline includes several steps: (i) tokenization: breaking text into meaningful units, (ii) tagging: assigning a part of speech tag to each token (noun, adjective, verb, etc.) depending on its usage in the sentence, (iii) parsing: extracting the dependency relationship between the words, and (iv) entity recognition: locating named entities in text and classifying them into categories. We added a PhraseMatcher class before entity recognition step to search for all possible phrases we defined. We converted the text into lowercase to allow for case-sensitivity. Abstracts containing both GOF and LOF acronyms were filtered out. We applied an iterative search on all available abstracts for each mutation. When there was a disagreement, i.e. a mutation was found as GOF in one abstract, and LOF in another abstract, it was labeled as “Mixed”, and excluded from the dataset. The remaining mutations were labeled as “GOF” or “LOF”, if one of the keywords was found in at least one associated abstract.

#### Selection of neutral variants

We used variants selected from the Genome Aggregation Database (gnomAD release 2.0.2) to compare GOF and LOF (function affecting) mutations which were very likely to be neutral variants ^27^. The gnomAD database provides harmonized exome and genome sequencing data from unrelated individuals sequenced as part of various disease-specific and population genetic studies. The individuals affected by severe disease and their first-degree relatives were not included in the dataset, allowing it to be used as a reference dataset that represents the general population. We randomly selected 10^6^ variants and extracted 134,450 likely neutral variants that occur in the same genes as our GOF and LOF mutations.

### Feature annotations

We used several tools and resources to annotate an extensive set of features (Table S1). This subsection is organized as (i) variant-level features, (ii) gene-level features, and (iii) protein-level features.

#### Annotation of variant-level features

All variants were first annotated with Ensembl Variant Effect Predictor (VEP, release 93) based on the Human GRCh37 assembly. Variant annotation tools provide a prediction for each transcript that a variant overlaps with. To filter one prediction result per variant, we selected the annotations of the Ensembl canonical transcripts ^28^. The canonical transcript for a human gene is defined by Ensembl based on following: (1) Longest CCDS translation with no stop codons; (2) If no (1), choose the longest Ensembl/Havana merged translation with no stop codons; (3) If no (2), choose the longest translation with no stop codons; (4) If no translation, choose the longest non-protein-coding transcript.

#### Consequence

The predicted variant consequence by Ensembl (i.e., missense, frameshift, stop_gain, splice, etc.) and the impact rating (i.e., high, moderate, low and modifier) indicating the severity of the consequence were used as two separate features ^28^.

The variant classes were defined by Ensembl as: (i) single nucleotide polymorphism (SNP), (b) insertion of one or several nucleotides (insertion), and (iii) deletion of one or several nucleotides (deletion).

#### Impact

The predicted functional impact of amino acid substitutions by SIFT and PolyPhen-2 were calculated by VEP ^16, 17, 28^. Both predicted terms and scores were used as features. The highest allele frequency (max_af) observed in any population from 1000 Genomes Project, Exome Sequencing Project (ESP) or gnomAD was calculated by VEP and used as a feature ^27, 29^. We used several available plugins with VEP (https://github.com/Ensembl/VEP_plugins) to extend functional impact annotations including CADD, Condel and REVEL. Combined Annotation-Dependent Depletion (CADD) scores have been widely used for predicting the deleteriousness of single nucleotide variants and insertion/deletion variants in the human genome^15^. Condel is a predicted deleteriousness score of missense variants calculated based on the pre-calculated SIFT and PolyPhen-2 scores from the Ensembl API ^16, 17, 30, 31^. REVEL is a score predicting the pathogenicity of missense variants ^18^.

### Position

The relative position of the mutation in the protein sequence, cDNA, and CDS were calculated by dividing the mutation position by the sequence length. The feature indicates the percentile interval in which the mutation occurs: (1): 0-10%, (2): 10-20%, (3): 20-30%, (4): 30-40%, (5): 40-50%, (6): 50-60%, (7): 60-70%, (8): 70-80%, (9): 80-90%, (10): 90-100%.

### Annotation of gene-level features

#### Essentiality

Genic intolerance was evaluated using different tools. We calculated LoFtool and ExACpLI scores using VEP plugins. LoFtool is a gene intolerance ranking score based on the ratio of LoF (stop-gain, splice site disrupting, frameshift) mutations to synonymous variants from the Exome Aggregation Consortium (ExAC) dataset (http://exac.broadinstitute.org/), but adjusted for the *de novo* mutation rate and protein evolutionary conservation ^32^. A lower LoFtool score indicates more intolerance to functional variation. The ExACpLI score represents the probability of a gene being loss-of-function intolerant (pLI) computed by the ExAC consortium based on observed and expected protein-truncating variant counts in exome sequence data for 60,706 individuals of diverse ethnicities ^33^.

The Residual Variation Intolerance Score (RVIS) is a gene-based score calculated based on common functional variation compared to neutral variation from whole exome sequence data from large datasets from the human population ^34^. While a positive RVIS score points to more common functional variation, a negative score indicates intolerance. We used the latest RVIS release (v4), which is based on the ExAC v2 (http://gnomad.broadinstitute.org/), the consensus coding sequence (CCDS) release 20 (http://www.ncbi.nlm.nih.gov/CCDS), and Ensembl release 87 ^34^.

The *de novo* excess rate (Z-score) estimates the rate of *de novo* mutation per gene and calculates if a given gene contains more *de novo* mutations than expected by chance alone ^35^. The predicted functional indispensability score (IS) estimates the essentiality based on a gene’s network and evolutionary properties ^36^.

The gene damage index (GDI) provides an accumulated mutational damage estimate (based on CADD scores) for each gene in the healthy human population from the 1,000 Genomes Project database ^15, 37, 38^.

Selective evolutionary pressure acting on each gene was estimated by the McDonald–Kreitman neutrality index at the population level ^39^. The neutrality index compares the numbers of non-silent and silent substitutions to the numbers of non-silent and silent polymorphisms, and measures the direction and degree of departure from neutrality.

The length of the coding sequence (CDS) of the canonical transcript and number of paralogs were extracted from Ensembl BioMart release 75 ^40, 41^.

#### Inheritance

Mode of inheritance (MOI) of each gene was extracted from the HGMD database and sub-categorized as autosomal recessive (AR), autosomal dominant (AD), autosomal recessive/dominant (ADAR), X-linked dominant (XLD), X-linked recessive (XLR), and unknown ^26^.

#### Pathogenicity

The mutation significance cutoff (MSC) introduces gene-level thresholds for predicted mutation impact scores such as CADD, PolyPhen-2 and SIFT ^15–17, 42^. In this study, we included MSC-CADD cutoffs with 95% confidence interval.

#### Annotation of protein-level features

Each protein can generate a number of alternative protein sequences (isoforms) through different biological events such as alternative splicing, alternative promoter or polyadenylation site usage, etc. The canonical protein isoform is defined as the most relevant sequence and/or the most similar sequence. To annotate the correct protein isoform, we used Ensembl protein IDs instead of UniProt IDs of canonical isoforms. As a result, in this study, we considered protein sequences that were translated from canonical transcripts but not necessarily canonical isoform protein sequences.

#### Amino acids

The reference and alternative amino acid types, as well as their positions were extracted by Ensembl VEP and used as features ^41^. When there was an insertion or deletion, only the first amino acid type and position were considered.

#### Protein domain

Pfam is a database of protein families where families are represented by multiple sequence alignments and hidden Markov models ^43^. Pfam protein domain information of all genes in our database were downloaded from the Ensembl BioMart (release 99) using Ensembl protein IDs with version corresponding to the canonical transcripts as annotated by Ensembl VEP ^40, 44^. We also searched the InterPro database for the proteins lacking Pfam domain information in Ensembl BioMart. When a protein exists in a protein database but the mutation position did not fall into a known protein domain, it was considered as “outside” of the domain, and when a protein was absent from both the Pfam and InterPro databases, it was considered as “unknown” domain.

Transmembrane helix regions were derived from Ensembl VEP, which uses predictions by TMHMM ^45^.

Post-translational modification (PTM) regions for phosphorylation, acetylation, ubiquitination, methylation, N-linked glycosylation, and O-linked glycosylation were downloaded from the dbPTM. The dbPTM is a database that includes experimentally validated PTM sites from available databases and manually curated PTMs from the literature ^46^.

#### Disordered region

Disordered regions are defined as regions that lack a stable well-defined tertiary structure under native conditions. IUPred2A was used to predict intrinsically disordered protein regions (IDPRs), and disordered binding regions ^47^. IUPred2A is based on pairwise energies estimated from observed amino acid compositions and returns probabilities of each residue being part of a disordered region and being part of a disordered binding region. We used the default (long) disorder prediction type predicting global structural disorder that encompasses at least 30 consecutive amino acid residues of the protein. Clark’s distance was used to estimate the amino acid composition complexity of each protein ^48, 49^ from a previously published study ^37^.

#### Solvent accessible area and secondary structure

NetsurfP-1.1 was used to predict the solvent accessible surface area and secondary structure for each protein ^50^. This method is based on neural networks that had been trained on experimentally determined protein structures. NetsurfP-1.1 returns solvent accessible surface area (ASA) given in Å^2^, relative surface area (RSA), RSA class ((buried or exposed i.e., below or above 25% exposure), reliability of each prediction (Z-fit). In addition, NetsurfP-1.1 provides the probability of each residue being in an alpha-helix, beta-strand, or coil region.

### Feature importance

To evaluate the discriminatory power of our features, we used (i) Fisher’s exact test, (ii) Boruta algorithm.

#### Fisher’s exact test

We used SciPy v1.1.0 python library to perform Fisher’s exact test on 2x2 contingency tables. Each category of a feature was evaluated separately based on its presence/absence in GOF and LOF datasets. All continuous features were converted to categorical features to be able to calculate p-values with Fisher’s exact test. When available, we used the original definitions of categorical features instead of continuous values. When there were no defined cutoff values, we identified three classes based on distributions of the continuous values. First, the interquartile range (IQR) of non-normally distributed data which is defined as IQR=Q3-Q1, where Q1 is the first quartile and Q3 is the third quartile, was calculated ^51^. Then, the boundaries for categories were identified as median±0.25*IQR. A drawback of converting continuous features into categorical variables is that it might result in loss of information and power. A full list of categorical features can be found in Table S1. All p-values were corrected using the Bonferroni correction method.

For the significant p-values showing associations between GOF versus LOF mutations, GOF versus neutral variants, and LOF versus neutral variants, we also calculated the logarithm of the odds ratio (OR) to compare features of GOF and LOF mutations by using neutral variants as background. That is, we determined log-odds ratios indicating enrichment for GOF mutations and LOF mutations separately, compared to neutral variants.

### Boruta algorithm

We used the Boruta feature selection package implemented in R, which is a wrapper algorithm built around the random forest classifier ^52^. Whereas traditional algorithms return a subset of features that gives the best possible classification result (minimal-optimal approach), the Boruta algorithm identifies all features that are somewhat relevant (weakly or strongly) for classification (all-relevant approach).

In random forest, the importance of a feature is calculated as the loss of accuracy of classification caused by the random permutation of feature values. It is computed separately for all trees in the forest. Then the Z-score is computed by dividing the average accuracy loss by its standard deviation. The Z-score cannot be directly used to measure feature importance as a reference value is needed. For this reason, the Boruta algorithm creates shadow features which are copies of the original features with randomly shuffled values. Then, the maximum importance of shadow features can be used as a reference to decide if original features are important. The features, which significantly outperform the best shadow feature, are interpreted as confirmed.

First, we omitted the features SIFT, PolyPhen-2, Condel and REVEL scores which are only available for missense mutations to deal with high missing rates in our database. We treated the missing categorical data as another category, “unknown”. The continuous features were used without binning. A full list of features can be found in Table S1. The remaining missing values in continuous features were imputed using the MICE R package with the predictive mean matching (pmm) imputation method ^53^. We also performed scaling on data to normalize the features.

To deal with imbalanced GOF and LOF dataset sizes, we used the ROSE R package with under-sampling option ^54^. The probability of the minority class (GOF) was chosen as p=0.4. In the resulting balanced dataset of 2,400 mutations, there were 1,254 GOF and 1,146 LOF mutations.

### Overrepresentation analysis

The Gene Ontology (GO) and protein class overrepresentation analysis were performed using PANTHER for gene sets having GOF mutations and LOF mutations separately compared to all human genes in the PANTHER database ^55–57^. Fisher’s exact test has been performed with Bonferroni correction.

### GOF/LOF Web-based Interface

The GOF/LOF web-based interface generated in this study allowing users to easily explore and download the data has been developed by R-Shiny platform (www.rstudio.com/shiny).

## Results

### Database of GOF and LOF mutations

Currently, there is no available genome-wide database of ascertained germline GOF and LOF disease-causing mutations in human. Such a database could serve as an invaluable resource with which to study features of mutations causing different diseases and to develop computational prediction tools. To fulfill this need, we developed an automated pipeline based on text-mining and NLP to extract all disease-causing GOF and LOF mutations from paper abstracts and established a comprehensive database of GOF and LOF mutations. We searched 123,737 paper abstracts associated with disease-causing mutations in the HGMD database (Professional version 2019.1) for different possible terms describing GOF and LOF mutations ^25, 26^. It should be noted that the terms defining GOF and LOF are inconsistent, with examples including “gain-of-function”, “gain of function”, “GoF”, “GOF”, making it challenging to analyze in an automated fashion. Therefore, we used NLP to search for different patterns and phrases matching the patterns we defined. The NLP framework helps to extract phrases with specific pattern types from a given text. We thus identified 1,237 mutations as GOF and 10,133 mutations as LOF. To the best of our knowledge, this is the largest GOF and LOF mutation database established to date. In Table S2, we provide a subset of the predicted GOF/LOF database containing 949 GOF and 8,592 LOF mutations from the publicly available version of HGMD December 2019. Although the majority of both GOF (89%) and LOF (55%) mutations in our database were missense mutations, the frameshift (23%) and stop-gains (19%) account for a significant percentage of LOF mutations. To address the question of whether the discriminative features we observed resulted from this difference, we compared the various features specifically in the proportion of missense GOF and LOF mutations. In our database, we have 1,101 missense GOF and 5,507 missense LOF mutations.

### Discriminative variant-level features of GOF and LOF mutations

#### LOF mutations have a higher impact on protein sequence and function

To evaluate the significance of the mutational effects on protein function, we compared the consequence (e.g., missense, stop-gain, frameshift), the impact rating (e.g., high, moderate, low), and the variant class (e.g., indel, SNV) of the GOF and LOF mutations and likely neutral variants from gnomAD by using the Ensembl Variant Effect Predictor (VEP) tool ^28^ (see Methods). GOF mutations were mostly predicted to be missense (89%), whereas LOF mutations were predicted to be missense, frameshift, and stop-gain (55%, 23%, and 19%, respectively) (Fig. 1a). The variants with high impact (i.e., frameshift, splice acceptor/donor, start-lost, stop-gained, stop-lost) and moderate impact (i.e., inframe insertion/deletion, missense) were found to be highly enriched among both GOF and LOF mutations as compared to neutral variants, while variants with low and modifier impacts (i.e., synonymous, intron) were more enriched among the neutral variants (Fig. 1b). Since we specifically selected the GOF and LOF mutations from the disease-causing mutations collected in the HGMD database, and the reference dataset comprises likely neutral variants from the gnomAD database, this result affirms the pathogenic effect of the GOF and LOF mutations in our database. More specifically, predicted functional consequences on protein sequence showed that LOF mutations have higher (disruptive) impact in the protein than GOF mutations as they often resulted from a frameshift or stop-gain. The missense, frameshift, and stop-gained mutations exhibited significant differences between the GOF and LOF mutation datasets (Fisher’s exact 2-sided, Bonferroni corrected P-values were 6.26×, 2.92×10^-71^, 4.83×, respectively) (Table S3).

We also analyzed the differences in variant classification of GOF and LOF mutations, and neutral variants. Whereas the GOF dataset had similar proportions of insertion, deletion and single nucleotide variations (SNVs) to neutral variants, the LOF dataset had fewer SNVs and more insertions and deletions. Comparing the variant class differences between GOF and LOF mutations by Fisher’s exact test indicates that GOF mutations are significantly enriched among SNVs (Bonferroni corrected 2-sided P-value=1.21×) whereas LOF mutations are enriched among micro-deletions and micro-insertions (P-values=8.94× and 1.0×) (Fig. 1c, Table S3). Overall, the impact rate, consequence for protein sequence, and variant class features of GOF and LOF mutations indicate that LOF mutations have a greater effect on protein sequence.

Using our GOF/LOF mutations database, we evaluated the predictive power of various computational tools designed to predict different aspects of mutation effect on function. We calculated the functional effect scores of the mutations by polymorphism phenotyping (PolyPhen-2), sorting intolerant from tolerant (SIFT), Combined Annotation Dependent Depletion (CADD), consensus deleteriousness (Condel), and rare exome variant ensemble learner (REVEL) tools ^15, 17, 18, 30, 58^. PolyPhen-2, SIFT, Condel, and REVEL are only available for missense mutations. Therefore, non-missense mutations could not to be scored by these tools. LOF mutations failed to receive a score more often compared to GOF mutations (P-values: 7.86×, 1.9×, 1.82×, 2.94×, respectively).

Remarkably, in the missense GOF/LOF database, missense LOF mutations were more often predicted to be “probably damaging” by PolyPhen-2, “deleterious” by Condel and SIFT, having high scores (≥20) by CADD, and “likely pathogenic” by REVEL (P-values: 1.13×, 1.06×, 3.77×, 2.3×, 0.001, respectively) as compared to missense GOF mutations. Overall, pathogenicity prediction methods tend to predict missense LOF mutations as more damaging than missense GOF mutations, whereas GOF mutations were more often predicted to be “benign” by PolyPhen-2, “neutral” by Condel, “tolerated” by SIFT, having medium scores (between 10-20) by CADD, and “benign” by REVEL (P-values: 6.28×, 4.14×,1.4×, 0.003, 0.0007, respectively). Fig. 2 shows the differences in score predictions for GOF, LOF and neutral mutations. A low specificity of SIFT and PolyPhen for predicting GOF and LOF missense mutations in *ABCC8*, *KCNJ11*, and *GCK* genes has been previously reported ^59^. Our results confirm that current bioinformatics tools are relatively successful at capturing LOF mutations (likely due to being trained mostly by LOF rather than GOF mutations), whilst GOF mutations were more often incorrectly predicted to be benign.

**Fig. 2.**
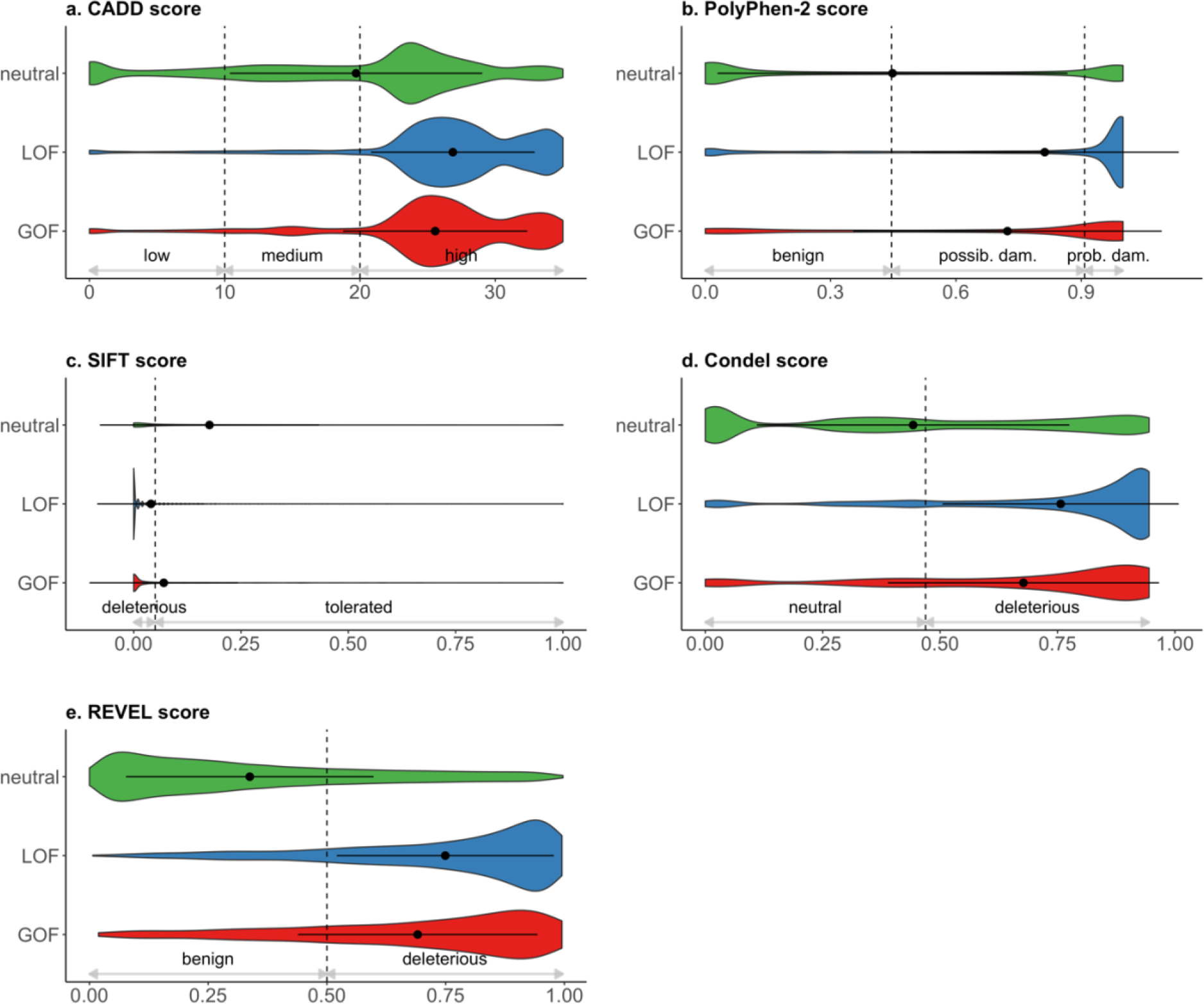
Pathogenicity prediction scores for GOF/LOF mutations by various methods. The x-axis corresponds to the predicted scores of GOF and LOF mutations, and likely neutral variants. The dashed lines represent the cutoff values we used for categorizing the scores (see Methods). **a)** The variants with higher CADD scores are predicted to be more deleterious. A CADD score of 10 indicates a variant to be among the 10% most deleterious mutations in the human genome, 20 indicates it to be among the top 1%, 30 indicates it to be among the top 0.1%, etc. **b)** The PolyPhen-2 score ranges from 0.0 (tolerated) to 1.0 (deleterious). The variants with less than a 0.446 score are predicted to be “benign”, whereas variants with more than a 0.908 score are predicted to be “probably damaging”. **c)** The SIFT score ranges from 0.0 (deleterious) to 1.0 (tolerated) where substitutions with a SIFT score < 0.05 are predicted to be “deleterious” and all others “tolerated”. **d)** The Condel score ranges from 0.0 (tolerated) to 1.0 (deleterious) where a substitution is predicted to be “deleterious” if the Condel score is greater than 0.469, and “neutral” otherwise. **e)** REVEL score ranges from 0 to 1 and variants with higher scores are predicted to be more likely to be pathogenic. REVEL scores above 0.5 indicate “likely disease causing” and scores below 0.5 indicate “likely benign” variants.

### Discriminative gene-level features of GOF and LOF mutations

In our database, we identified 135 genes having both GOF and LOF mutations, 1,437 genes having only LOF mutations, and 154 genes having only GOF mutations. In the following analysis, we compared the 289 genes harboring GOF mutations to 1,572 genes harboring LOF mutations.

#### GOF mutations tend to occur in essential genes

A gene is defined as essential when it does not tolerate loss-of-function mutation. In other words, when an essential gene loses its function, it results in lethality or a profound loss of fitness ^60^. The genes with GOF mutations in our database were predicted to be significantly more intolerant compared to the genes with LOF mutations by all five tools (Bonferroni-corrected 1-sided P-values are 6.78×, 1.83×, 3.18×, 2.3×, and 0.001 by Z-score (*de novo* excess rate), RVIS, LoFtool, IS (predicted indispensability score), and ExACpLI, respectively) (Fig. S1, Table S3). Z-score and RVIS evaluate functional variation; LoFtool and ExACpLI calculate ratio of protein truncating mutations; and indispensability score (IS) combines network and evolutionary properties (see Methods for details). In addition, when compared to neutral variants, LOF mutations occur significantly less frequently in intolerant genes (Table S6). Fig. 3 illustrates the differences in gene essentiality between GOF, LOF and neutral variants based on different metrics. The essentiality of the genes also remained significantly different for only missense GOF and LOF mutations (Fig. S1).

**Fig. 3.**
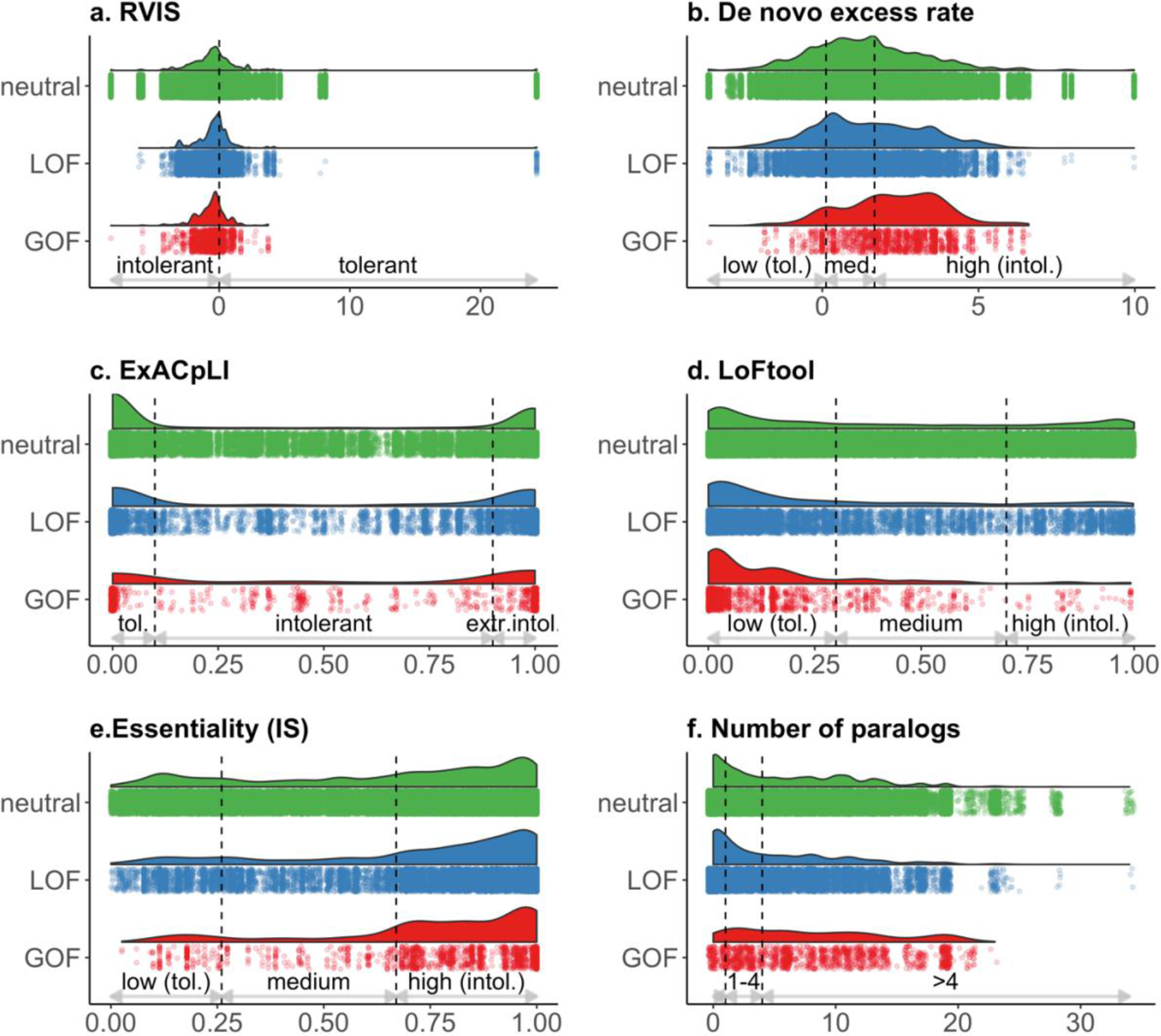
Essentiality predictions of the genes with GOF and LOF mutations, and likely neutral variants by various methods. In panels a-e, the x-axis corresponds to the predicted essentiality scores of the genes in each dataset, whereas in panel f the x-axis corresponds to the number of paralogs of the genes. The dashed lines represent the cutoff values we used for categorizing the features. When there were no defined cutoff values, three classes were identified based on distributions of the values (see Methods). **a)** A positive RVIS score is indicative of more common functional variation, a negative score implies intolerance. **b)** *de novo* excess rate (Z-core) is an estimation of the *de novo* mutation rate and a higher rate indicates intolerance. **c)** ExACpLI score represents the probability of a gene being LOF intolerant. **d)** a lower LoFtool score indicates more intolerance to functional variation. **e)** Predicted indispensability score indicates the essentiality based on a gene’s network and evolutionary properties. **f)** Number of paralogs were shown in three categories: having more than 4 paralogs, 1-4 paralogs, and being a singleton.

Paralogs are genes that have been duplicated (or even amplified) within the same organism during evolution, sharing a common ancestor. Paralogous genes are expected to be more tolerant to mutations since they may still exhibit some functional redundancy and hence can compensate to some extent for loss of function of one paralog ^61–64^. The degree of protection afforded also depends upon divergence in tissue distribution and molecular functionality over evolutionary time. When paralogs maintain related functions, they can buffer each other’s loss, as inactivation of one copy may not affect the other(s) ^64–66^. However, when paralogs have different or interdependent functions, they tend to be more intolerant, i.e. less protective against deleterious mutations ^66^.

To relate paralogy to GOF and LOF mutations, we determined the number of paralogs of each gene by Ensembl BioMart ^44^. This showed that GOF mutations occur more frequently in genes that have more than 4 paralogs (67%) as compared to LOF (41%) and neutral mutations (50%) (Figs. 3f and 4a). Loss of function in the genes that have more than 4 paralogs appears to be buffered by the other paralogs. The genes with GOF mutations more often had more than four paralogs compared to the genes with LOF mutations (P-value=5.08×), and the genes with LOF mutations were found disproportionally in singleton genes compared to GOF mutations (P-value=1.02×) (Table S3).

**Fig. 4.**
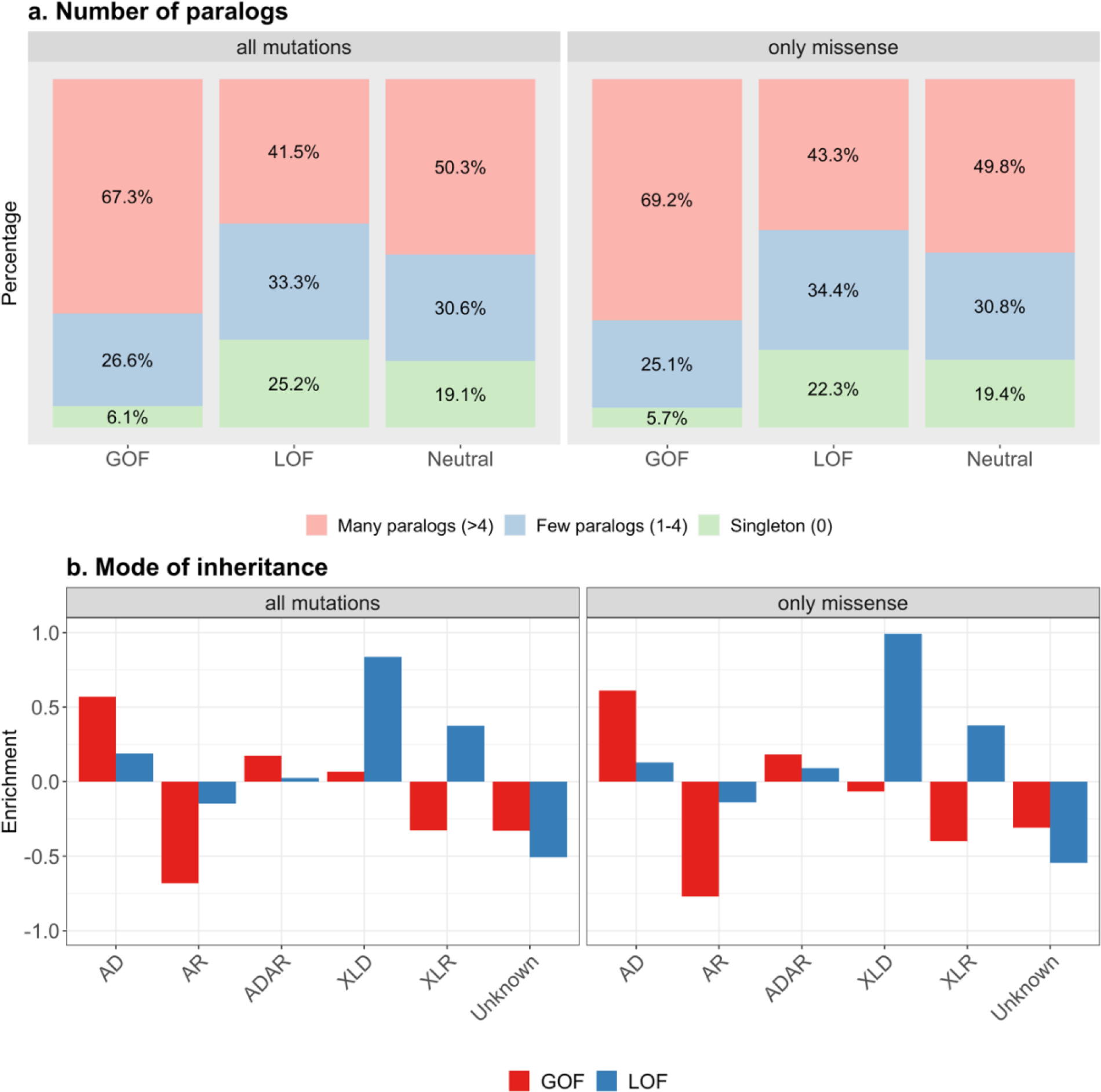
Number of paralogs and mode of inheritance. **a)** Fraction of GOF and LOF mutations, and likely neutral variants in genes with different numbers of paralogs. The y-axis corresponds to having more than 4 paralogs (many), having 1-4 paralogs (few), and being a singleton (none). The left-hand panel shows the fractions in the whole dataset, the right-hand panel shows the fractions in the subset of missense mutations. **b)** Mode of inheritance of the genes with GOF and LOF mutations. The y-axis corresponds to the log10 of the odds ratio of the enrichment of inheritance type in GOF and LOF mutations compared to likely neutral variants. The x-axis corresponds to autosomal dominant (AD), autosomal recessive (AR), autosomal dominant and recessive (ADAR), X-linked dominant (XLD), X-linked recessive (XLR), and not known (Unknown). Positive and negative values correspond to overrepresentation and underrepresentation in GOF/LOF mutations as compared to neutral variants, respectively. The left-hand panel shows the fractions in the whole dataset, the right-hand panel shows the fractions in the subset of missense mutations.

As genetic variant intolerance metrics are often correlated with coding sequence (CDS) length, we investigated the differences in CDS by defining 3 categories (long, medium and short) based on the distribution of sequence length in our database (see Methods) ^60^. We identified significant enrichment for longer length (>1,747 base pairs) in the genes with GOF mutations (P-value=5.19×) and medium length (592-1,747 base pairs) in the genes with LOF mutations (P-value=1.06×) suggesting that GOF mutations occur significantly more frequently in genes with longer CDS. The number of paralogs and the coding sequence length of genes were also significant features to distinguish missense GOF and LOF mutations. Missense GOF mutations occur more frequently in the genes that have paralogs (Fig. 4a), whilst the CDS was significantly longer (>1,747 base pairs) for missense GOF mutations (P-value=1.03×), and having medium length (592-1,747 base pairs) for LOF mutations (P-value=1.68×).

### GOF mutations occur mostly in genes underlying autosomal dominant conditions

Since loss-of-function mutations are often associated with autosomal recessive (AR) diseases, and autosomal dominant (AD) diseases are associated with either loss- or gain-of-function mutations, we analyzed the mode of inheritance (MOI) of the genes in our database ^67–69^. We found that the mutations in genes with autosomal dominant conditions were significantly enriched for GOF mutations (P-value=4.55×), whilst the mutations in genes with autosomal recessive conditions were enriched for LOF mutations (P-value=6.74×). Genes underlying X-linked dominant or X-linked recessive conditions were enriched for LOF mutations (P-values: 9.23×, and 1.06×, respectively). Moreover, compared to neutral mutations, GOF mutations were enriched in genes underlying autosomal dominant conditions, whilst LOF mutations were enriched in genes underlying X-linked conditions (Fig. 4b, Tables S5, S6). Overall, these results reveal the significant differences in the MOI of the disease genes harboring GOF and LOF mutations, consistent with the known association between mode of inheritance and mutation type70-72.

For the subgroup of missense GOF and LOF mutations, the MOI of the genes was the most significant difference; missense GOF mutations were significantly enriched in genes with AD inheritance (P-value=4.88×), whereas missense LOF mutations were significantly enriched in genes with AR, X-linked dominant, and X-linked recessive inheritance (P-values: 2.37×, 4.29×, and 8.07×, respectively) (Fig. 4b, Table S4).

### Discriminative protein-level features of GOF and LOF mutations

#### GOF and LOF mutations are enriched in certain amino acid substitution types

Each amino acid has distinct physicochemical characteristics that affect its role in protein structure and function. We analyzed the amino acid substitution spectra of GOF and LOF mutations first in our full database, and neutral variants from gnomAD, and then in the subset of missense mutations. In line with predicted mutational consequences, LOF mutations were significantly enriched for premature termination of the protein and frameshift. Interestingly, hydrophobic amino acids Valine (V), Methionine (M), Phenylalanine (F), Glycine (G), and Isoleucine (I) (P-values: 4.32×, 7.4×, 6.5×, 0.001, 0.01, respectively), as well as charged amino acids Lysine (K) (P-value=1.87×) and Glutamic acid (E) (P-value=0.03) were found to be enriched in alternate (i.e., substituting) residues in GOF mutations (Table S3, Fig. 5a).

**Fig. 5.**
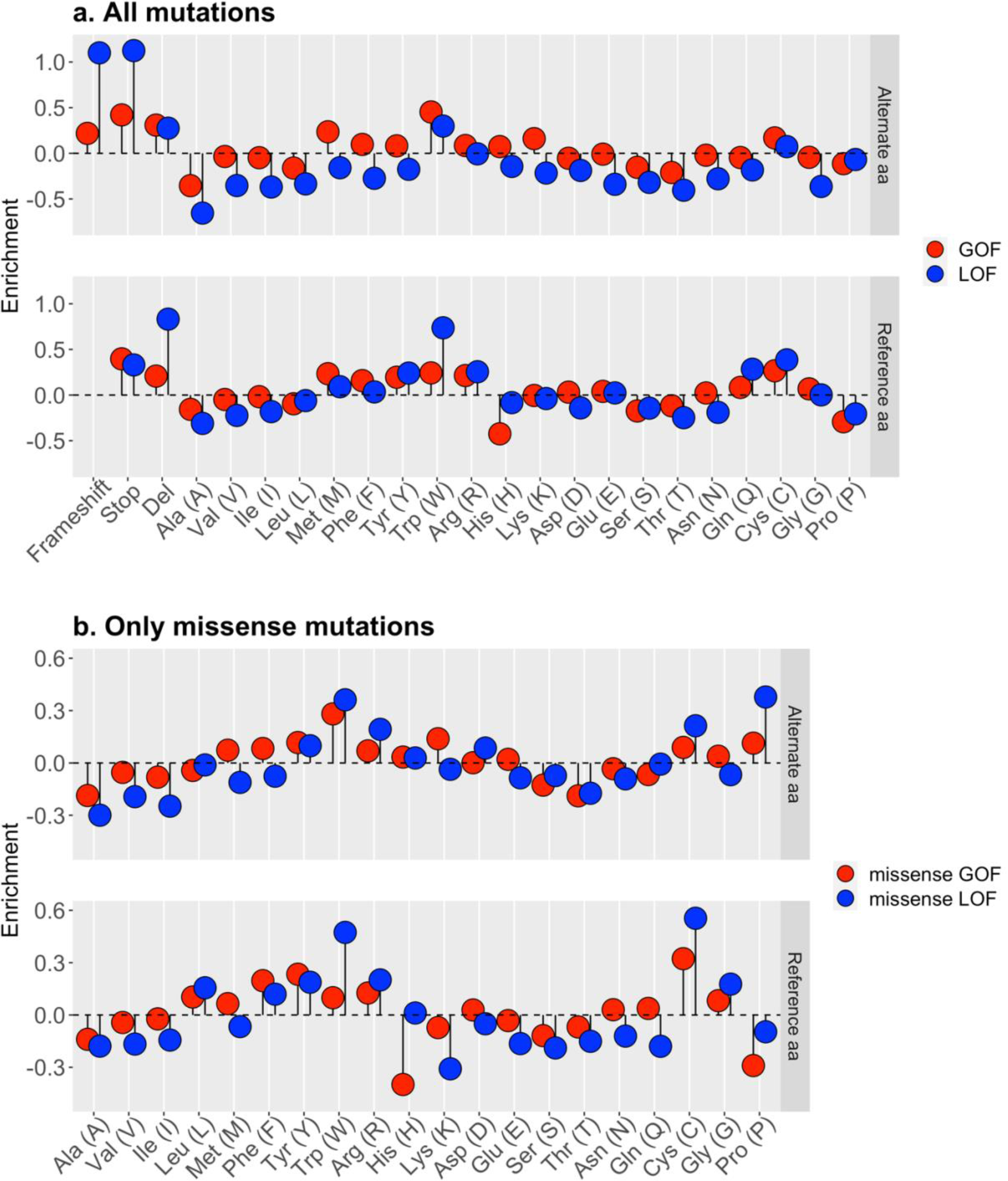
Amino acid changes in GOF and LOF mutations. **a)** Reference and alternate amino acid type enrichment of GOF and LOF mutations compared to likely neutral variants. The y-axis corresponds to the log10 of the odds ratio of the enrichment of reference amino acid type. The x-axis corresponds to 20 amino acid types plus frameshift, stop-gain (stop), and deletion (del). **b)** Reference and alternate amino acid type enrichment of missense GOF and missense LOF mutations compared to likely neutral missense variants. The y-axis corresponds to the log10 of odds ratio of the enrichment of reference amino acid type. The x-axis corresponds to 20 amino acid types. Amino acid types are arranged based on their physicochemical properties (e.g., hydrophobicity, charge).

Tryptophan (W) in the reference position was enriched for LOF mutations (P-value=5.2×). Furthermore, compared to neutral mutations, we observe mutations to Tryptophan (W) more frequently in both GOF (P-value=6.19×) and LOF mutations (P-value=1.05×), whereas mutations to Alanine (A) are less frequent in both GOF (P-value=9.3×) and LOF (P-value=1.88×) (Fig. 5a, Tables S4 and S5). Mutations to Tryptophan are highly associated with disease causation because Tryptophan is a hydrophobic and bulky amino acid typically found in the protein core; thus, mutation of a tryptophan residue, whether on the surface or buried, is likely to disrupt protein folding. Conversely, mutation to Alanine is less likely to be disease-causing since Alanine is a neutral, small amino acid and its introduction is less likely to have a dramatic effect on protein structure.

Missense GOF and LOF mutations also exhibited significant differences in terms of the types of amino acid involved (Fig. 5b). Among the 20 amino acid types in alternate (i.e., substituting) and reference (i.e., substituted) positions, only Proline (Pro/P) in alternate position was significantly more enriched (P-value=0.02) in missense LOF mutations compared to missense GOF mutations. Proline, which consists of a rotationally constrained rigid ring structure, typically breaks or kinks a helical structure, causing dramatic effects on protein structure.

However, an amino acid substitution matrix (20x20) showed significant differences for multiple substitution types (Fig. 6a). Remarkably, amino acid changes in GOF usually involve only subtle changes in the physicochemical properties of the mutated amino acid, and hence are less likely to cause misfolding and complete loss of function. For example, we observe frequent transitions from Isoleucine, Leucine, and Methionine to Valine (P-values: 1.8×, 0.008, 0.02, respectively) (all hydrophobic). Other significant transitions in GOF mutations were from Alanine (small amino acid) to Glycine (small amino acid), from Lysine (charged) to Glutamic acid (charged), from Glutamine (neutral polar) to Arginine (neutral polar), and from Phenylalanine (aromatic) to Tyrosine (aromatic). A substitution by Glycine is less likely to cause a drastic change since it is the simplest amino acid that possesses a single hydrogen atom as its side chain. By contrast, LOF mutations were associated with significant changes in the physicochemical properties of the amino acid. For example, transitions from the neutral Cysteine, Glycine and Leucine to the basic Arginine were more significantly enriched (P-values: 0.001, 0.014, 0.01, respectively). Cysteine is often involved in disulfide linkages that enhance protein stability. In addition, another substitution with significant structural effect in LOF mutations was from Leucine and Serine to Proline, which serves to break alpha helices.

**Fig. 6.**
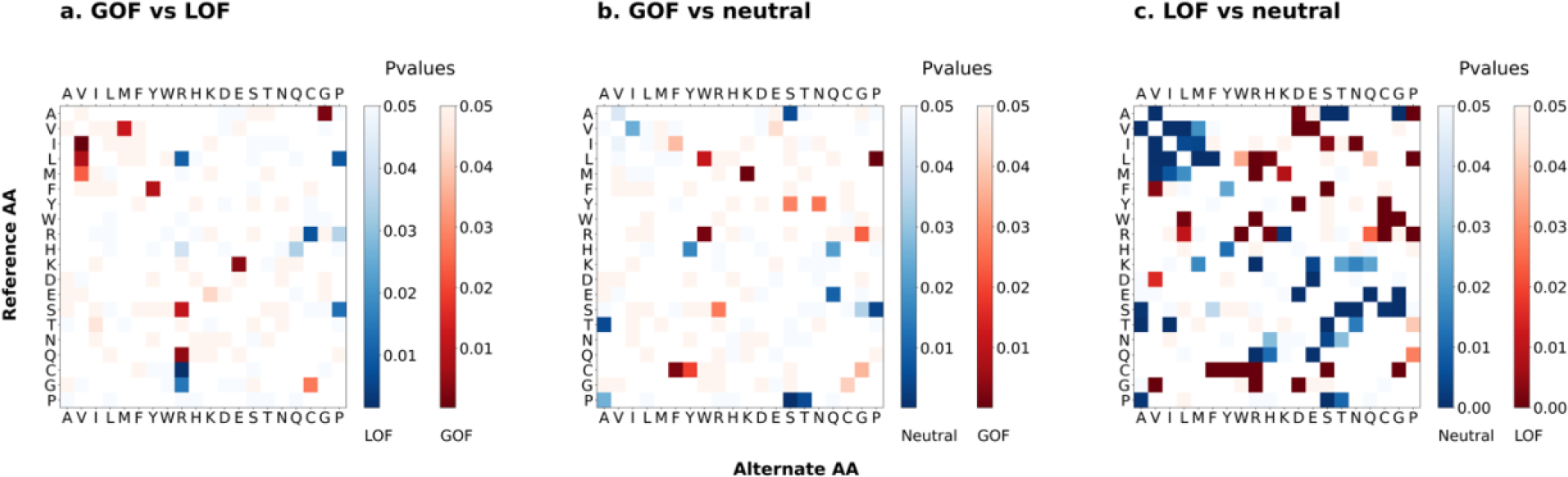
Enrichment of amino acid substitutions of missense mutations. **a)** Enrichment of amino acid substitutions between missense GOF and LOF mutations. Red color represents the substitutions more frequently seen in missense GOF mutations and blue color represents the substitutions more frequently seen in missense LOF mutations. **b)** Enrichment of amino acid substitutions between missense GOF mutations and likely neutral missense variants. **c)** Enrichment of amino acid substitutions between missense LOF mutations and likely neutral missense variants. Darker shade indicates more significant difference (i.e., lower P-values). The y-axis corresponds to 20 amino acid types in the reference protein sequence (i.e., substituted amino acid), and x-axis corresponds to 20 amino acid types in the alternate protein sequence (i.e., substituting amino acid). The p-values were calculated based on Fisher’s exact test.

#### Mutations at the beginning of sequences result in LOF

The relative position of the mutation in CDS, protein and cDNA provides information on the position of the mutation relative to the length of the sequence in question. LOF mutations were significantly more enriched in the first decile (1/10) of the CDS and protein sequences, and in the second decile (2/10) of the cDNA sequences compared to GOF mutations (P-values: 0.004, 0.004 and 0.02, respectively). This result is in line with the fact that frameshift and nonsense mutations that occur earlier in the sequence are more likely to result in misfolded protein which is unlikely to be functional.

Interestingly, missense LOF mutations were significantly enriched only in the last decile (10/10) of the cDNA sequences compared to missense GOF mutations (P-value=0.03).

#### GOF mutations occur in specific protein domains

The modularity of proteins is critical for protein function and may also have clinical relevance. For example, it has been previously suggested that GOF mutations often occur in specific protein regions including protein–protein interaction interfaces and allosteric regions ^69^. To further assess the effect of GOF and LOF on protein function, we used Pfam domains to examine if GOF and LOF mutations occur in particular domains, which are associated with specific functions ^43^. Interestingly, GOF mutations were found to be significantly enriched in various Pfam domains including STAT_alpha, ion_trans, STAT_bind, sod_cu (superoxide dismutase, copper/zinc binding domain), tropomyosin, vWA (von Willebrand factor type A), and SH2, which often mediates protein-protein interactions in signaling (P-values: 3.49×, 4.62×, 1.47×, 9.78×, 5.98×, 2.46×, 3.3×, respectively). The only Pfam domains found to be enriched for LOF mutations were p450 that are associated with the oxidation-reduction process, BRCT (BRCA1 C terminus), and Melibiase (P-values: 3.16×, 0.006, 0.002, respectively). Missense GOF mutations were found significantly enriched in the same Pfam domains (i.e., STAT_alpha, ion_trans, STAT_bind, sod_cu, with P-values: 4.12×, 2.79×, 4.86×, 2.08×, respectively).

GOF mutations were found disproportionally in transmembrane regions (P-val=1.3×) compared to LOF mutations. This enrichment disappeared when we considered only missense GOF/LOF mutations, demonstrating that the difference between GOF and LOF mutations is highly likely to be driven by protein truncating LOF mutations (i.e., frameshift, stop-gain, stop-loss). The proteins having GOF mutations were predicted to have lower Clark’s distance (P-value=1.54×), a term of reference to proteins with an amino acid composition similar to that of the average human protein (unbiased composition) ^37, 48, 49^. Low complexity regions are composed of a subset of amino acids (biased composition), and hence have greater distance from the average protein. Additionally, low complexity regions tend to be disordered. Our results are consistent with the notion that truncating LOF mutations are enriched in low-complexity disordered regions which often play a key role in signaling and regulation ^68, 69, 73^.

#### Missense LOF mutations are enriched in protein core regions

Predicted relative solvent accessibility (RSA) of proteins was found to be one of the most discriminatory features for missense GOF and LOF mutations although it did not show a significant difference when considering all mutations. Missense LOF mutations were enriched in buried regions, likely disrupting the stability and folding of the protein, whereas missense GOF mutations were found in regions more exposed to the solvent (P-value=5.33×), which likely have functional effect. Finally, missense GOF mutations were found to be enriched in disordered protein regions (P-value=3.88×). Disordered regions are considered to be evolutionarily less conserved and therefore might have higher tolerance to LOF mutations.

### Feature importance

To evaluate the discriminatory predictive power of our features, we used two different types of feature selection methods: (i) Fisher’s exact test, which is a statistical significance test calculating the association between two groups; and (ii) the Boruta algorithm, which is a wrapper built around the random forest machine learning classification algorithm ^52^. Fisher’s exact test compares two groups (GOF and LOF mutation datasets in our case) and assesses the correlation or dependence between the groups. Then, the important features can be selected by ranking the P-values based on a user-defined threshold. However, with the Fisher’s exact test, the interactions and correlations between feature variables or feature and target variables are not considered. Wrapper methods on the other hand measure the importance of features based on the classifier performance by iteratively selecting different feature subsets and evaluating the importance of each feature. Therefore, to evaluate the discriminative power of features based on the random forest classifier, as well as Fisher’s exact method, we also applied the Boruta algorithm.

We annotated 33 features (12 protein-level, 11 gene-level, and 10 variant-level). After binning the continuous features and binarizing the categorical features, 33 features were broken into 1,725 variables. We found 89 variables of 27 features to be significantly enriched by the Fisher’s exact test (2-sided Bonferronni-corrected pvalue<0.05) (Fig. 7a).

**Fig. 7.**
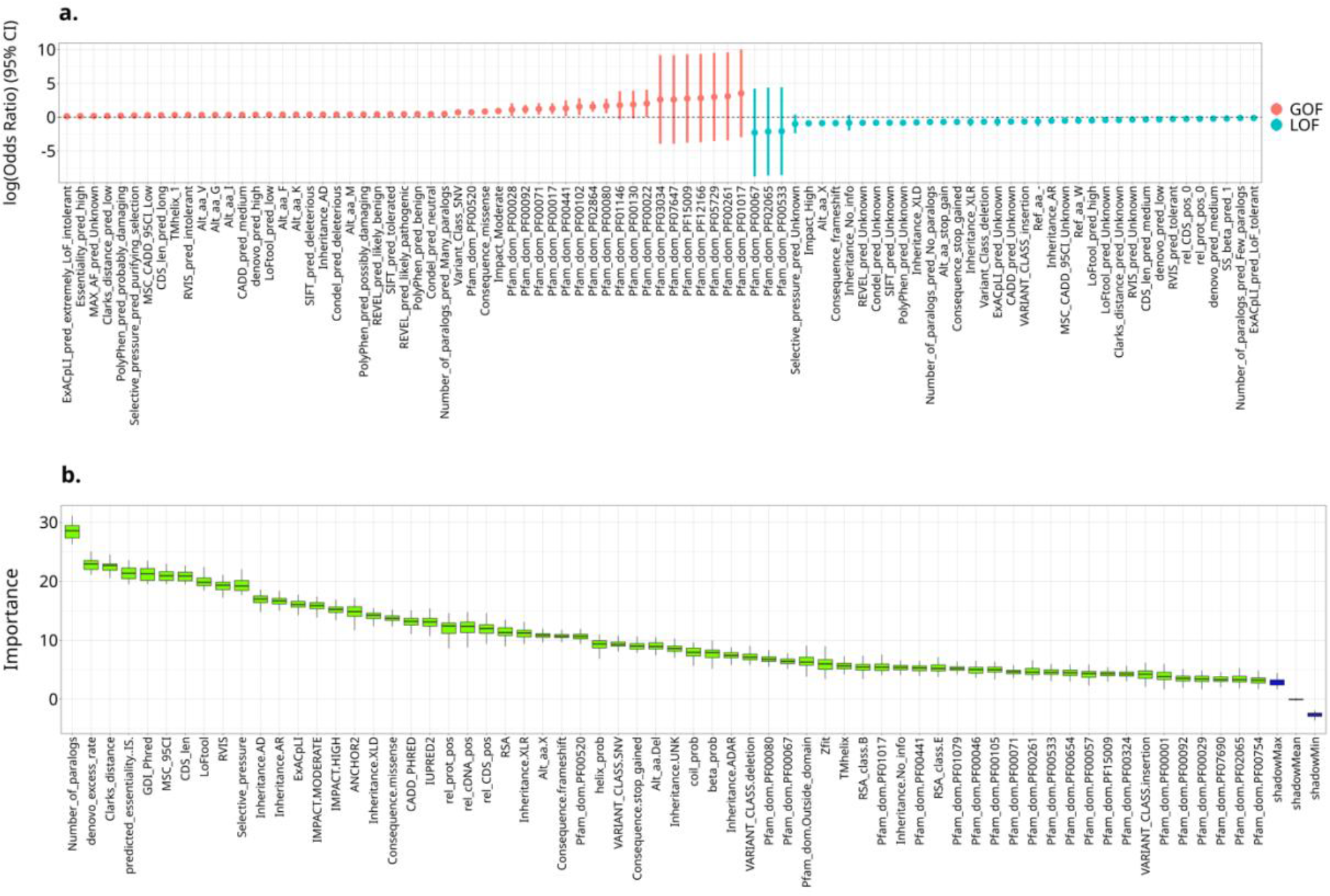
Enrichment of GOF and LOF mutations with gene- and protein-level features. **a)** Significant associations (Bonferroni-corrected 2-sided p-value < 0.05) calculated by Fisher’s exact test are shown. The y-axis corresponds to log10 odds ratio. Positive values indicate a feature’s enrichment for GOF mutations (shown in red) and negative values indicate a feature’s enrichment in LOF mutations (shown in blue). See Table S1 for details of the features. **b)** Feature importance ranked by the Boruta algorithm. The y-axis represents the Z-score of every feature in the shuffled dataset. The blue boxplots correspond to minimal, average and maximum Z-scores of shadow attributes (permuted copies of original features). The green boxplots represent Z-scores of confirmed attributes. A feature is rejected if it has a lower Z-score than that of max Z-score (if it under-performs the best shadow feature), and confirmed if it outperforms the best shadow feature.

When using the Boruta algorithm, SIFT, PolyPhen-2, Condel and REVEL scores were excluded from the feature list since these are not available for non-missense mutations. The missing values in the remaining features were imputed (see Methods, Figs. S2, S3). The categorical features were encoded by one-hot-encoding and the continuous features were used without binning. A balanced subset of 2,400 GOF and LOF mutations was randomly selected by using an under-sampling method ^54^. We determined 64 variables of 28 features as important (having a Z-score higher than the shadow features) with the Boruta algorithm (Fig. 7b).

The most important feature identified by the Boruta algorithm was the number of paralogs of a gene which is a metric for essentiality (Fig. 7b). Other important features include gene-level features associated with essentiality (*de novo* excess rate, IS essentiality score, gene damage index (GDI), CDS length, mutation significance cutoff (MSC), LoFtool, RVIS, selective pressure, ExACpLI) as well as MOI; protein-level features associated with protein sequence (Clark’s distance, disordered binding region, disordered region, protein domain, secondary structure); and variant-level features including impact rank, consequence type, CADD score, and variant class. Most of these features were also identified as important by Fisher’s exact test. Interestingly, the gene-level threshold values calculated by CADD scores (MSC-CADD) as well as GDI which measures of the mutational damage that has accumulated in the general population were also among the important features. Both GDI and MSC metrics correlate with gene tolerance/intolerance. The different results between Fisher’s exact test and the Boruta algorithm could be due to conversion of continuous variables into categorical features which leads to loss of information. Another important difference between these two approaches is that with Fisher’s exact test the correlations are evaluated independently, whereas the Boruta algorithm considers the interactions among the feature variables.

### Functions of genes significantly differ for GOF and LOF mutations

To analyze the functional properties of genes, we performed gene ontology and protein class overrepresentation analysis. The genes with GOF mutations were significantly enriched in metal ion, cation and ion transport among GO biological processes and in molecular functions related to ion channel and transmembrane transporter activity (Table S7). Transporter protein class was enriched in both gene sets, whilst metabolite interconversion enzyme and defense/immunity protein classes were enriched in the genes harboring LOF mutations. The transporter protein class involves the ion channels in which an altered channel activity may cause gain or loss of ion channel function.

## Discussion

We have generated and made available a comprehensive database of annotated GOF and LOF mutations. Our aim was to analyze gene-level and protein-level features of GOF and LOF mutations and to identify potential discriminative features, then to discuss the biological implications of these features in terms of how they contribute to protein gain- and loss-of function. We applied an NLP algorithm to the available abstracts of disease-causing mutations in the HGMD database followed by multiple annotation algorithms based on genomic and protein positions and sequence information. Our study yielded multiple discriminative features of GOF and LOF mutations suggesting a computational method based on machine learning techniques to classify the functional consequences of mutations.

To minimize the numbers of potential true positives and false negatives in our automated GOF/LOF prediction process, we excluded the abstracts mentioning both GOF and LOF mutations, and applied an iterative search of all available abstracts for each mutation and used the common prediction as label (GOF or LOF). Another remaining limitation of our approach was searching only paper abstracts instead of full text which is a more complex undertaking when applying NLP and requires free access. Whilst our study provides a first comprehensive GOF/LOF database, the data should be interpreted with caution as the GOF and LOF predictions are not manually curated.

To be able to apply Fisher’s exact test, we converted the continuous variables to categorical data. However, this approach might cause loss of information and variability in the data. To assess our findings, we also applied a machine learning based algorithm in which all variables were kept continuous. To compensate for missing values in our database, we applied data imputation which might itself introduce bias.

Among 33 tested features (1,725 variables), 27 features (89 variables) were identified as being significantly discriminative for GOF and LOF mutations with Fisher’s exact test, and 28 features (64 variables) with random forest-based algorithm (Fig. 7). Overall, 21 features were found to be significant with both approaches. For example, GOF mutations were enriched in some specific protein domains such as binding regions and domain interfaces where they may change the protein activation, allostery and interactions, whereas missense LOF mutations were enriched in protein core regions where they might disrupt the stability and folding of the protein. Overall, these results extend our knowledge of different properties of GOF and LOF mutations, and offer a powerful resource for analyzing these unique features. The findings of this study should improve our understanding of how GOF and LOF mutations impact protein function and ultimately cause disease.

## Supplemental Data

Supplemental Data include 3 figures and 7 tables.

## Acknowledgements

We thank Josue Barnes for his suggestions on available resources for protein-level feature annotations.

## Web Resources

HGMD, http://www.hgmd.cf.ac.uk/ac/index.php

gnomAD, http://gnomad.broadinstitute.org

Ensembl VEP, https://useast.ensembl.org/info/docs/tools/vep/index.html

VEP plugins, https://github.com/Ensembl/VEP_plugins

## Data availability

GOF and LOF mutations database and the annotated features used in the study are freely available at https://itanlab.shinyapps.io/goflof/.

## Author contributions

Y.I., A.S., A.P., S.B-D., and S.O. contributed to study conception, Y.I. and A.S. jointly supervised the study. C.S.B. organized the study, collected and structured the data, and performed the analyses. A.J. performed the variant annotations. D.S. performed the feature selection analysis and assisted the visualization. K.C. provided guidance on interpretation of the data. D.N.C. and P.D.S. provided the resources (HGMD dataset). G.N.N. provided guidance on machine learning. T.V.V. provided guidance on natural language processing. C.S.B., A.S., and Y.I. wrote the original draft. C.S.B., A.J., D.S., K.C., G.N.N., T.V.V., A.P., S.B-D., S.O., P.D.S., D.N.C., A.S., and Y.I. reviewed and edited the final manuscript.

