## Supplementary Figures for "Identification of Discriminative Gene-level and Protein-level Features Associated with Gain-of-Function and Loss-of-Function Mutations"

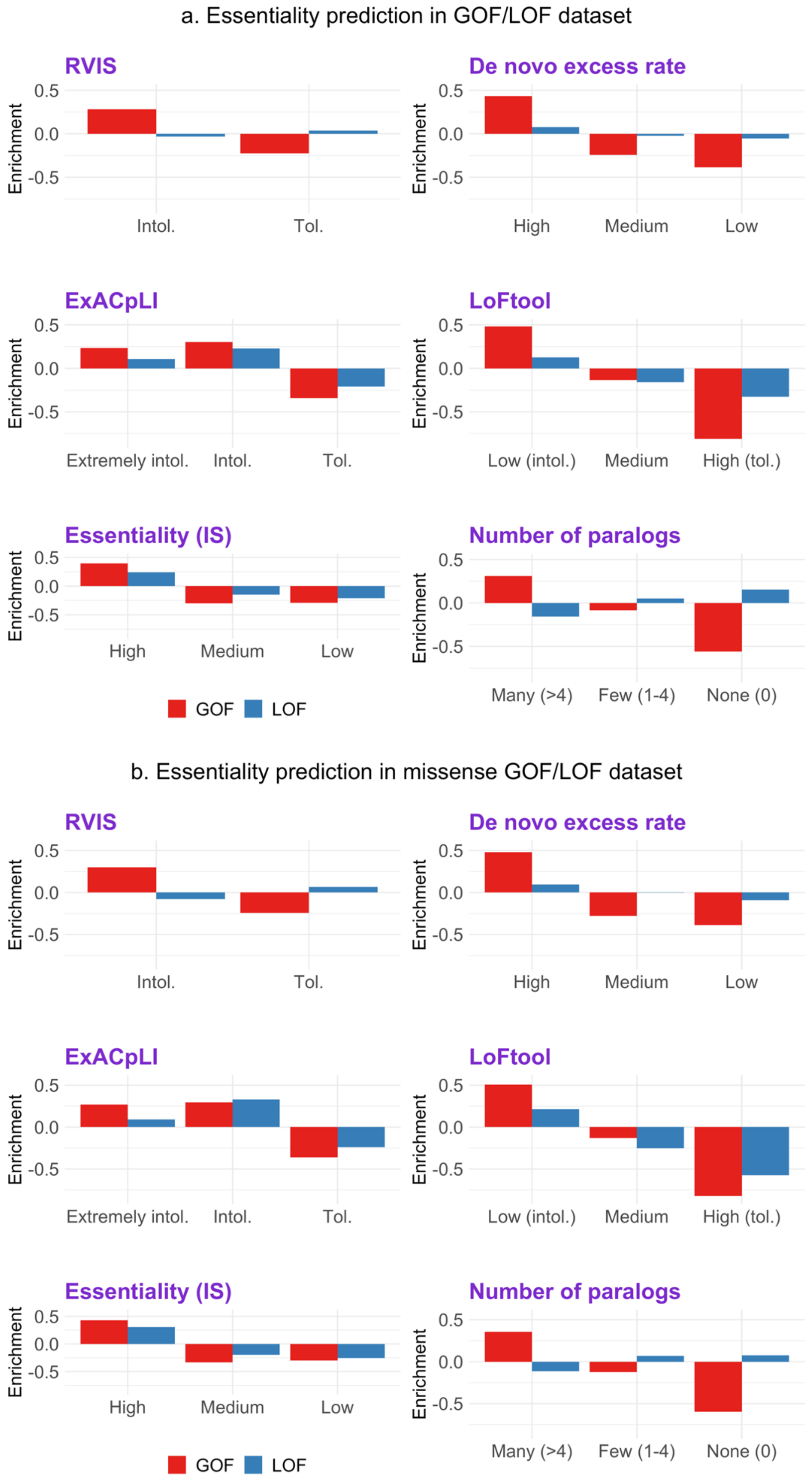
a. Essentiality prediction in GOF/LOF dataset
RVIS
De novo excess rate
ExACpLI
LoFtool
Essentiality (IS)
Number of paralogs
Enrichment
0.5
0.0
-0.5
Intol.
Tol.
High
Medium
Low
Extremely intol.
Low (intol.)
High (tol.)
Many (>4)
Few (1-4)
None (0)
GOF
LOF
b. Essentiality prediction in missense GOF/LOF dataset

**Figure S1.** Essentiality predictions of the genes with GOF and LOF variants by various methods. a) Essentiality enrichment in all GOF and LOF mutations compared to likely neutral mutations. b) Essentiality enrichment in only missense GOF and LOF mutations compared to likely neutral missense mutations.

The y-axis corresponds to the log10 of odds ratio of the enrichment of predicted scores in GOF and LOF mutations compared to neutral variants. Positive values correspond to the enrichment in missense GOF/LOF mutations compared to missense neutral variants. The x-axis corresponds to the predicted categories. To define the categories, we used the cutoff values given by the tool when available. When the cutoff values were not specified, we calculated two values based on interquartile range of distributions (see Methods for details).

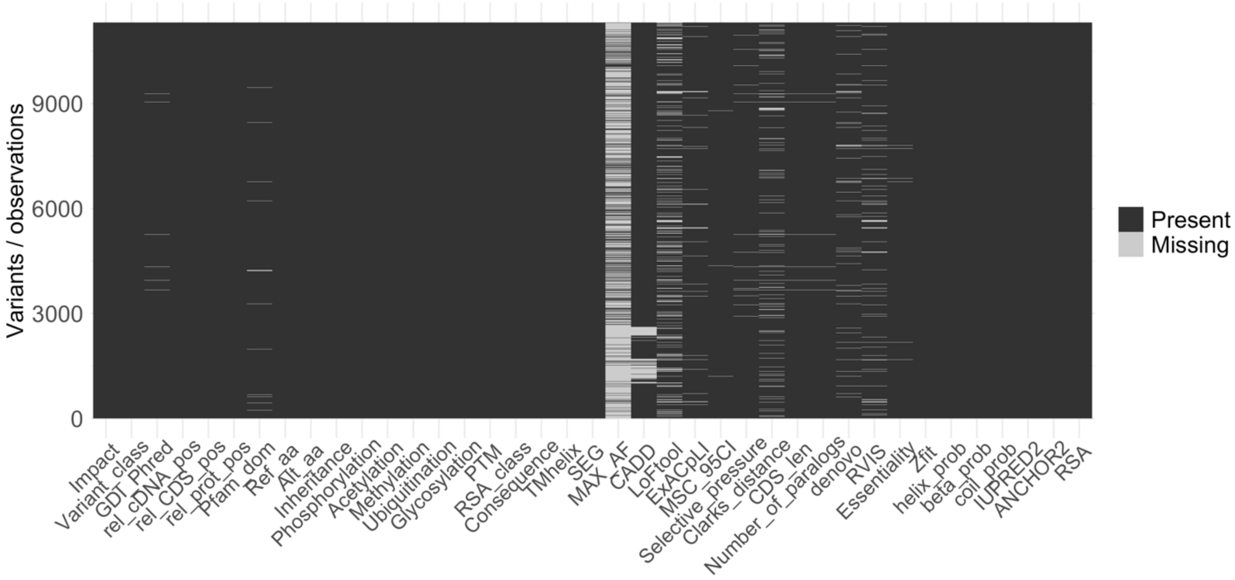


**Figure S2.** Missing values in GOF/LOF features before imputation. The x-axis corresponds to features, and the y-axis corresponds to the 11,370 GOF/LOF mutations. The light gray lines show the missing values in each feature.

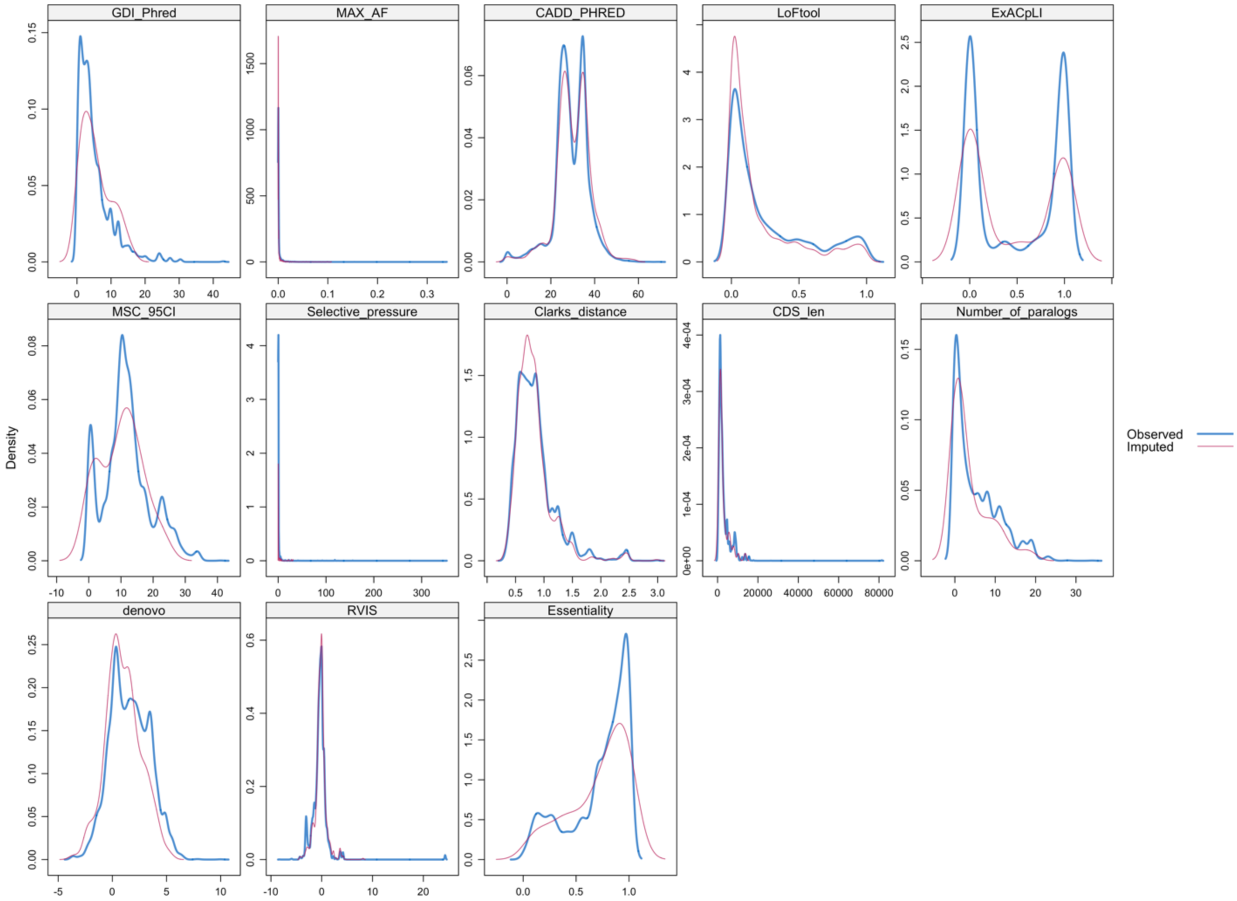


**Figure S3.** Density plots of imputed and observed features. All numeric features with missing data (see Fig S2) were imputed using R MICE package. The density of the imputed data is shown in magenta and the density of the observed data is shown in blue.
